# Investigating the role of CpG island DNA methylation at 3’UTRs in cancer

**DOI:** 10.1101/2024.10.18.619008

**Authors:** Claire Wilson, Aditi Kanhere

## Abstract

DNA methylation is one of the most important epigenetic processes that regulates gene expression. While the human genome is predominately methylated, CpG-rich regions known as CpG islands (CGIs) are unmethylated and are sites of transcriptional control. While the importance of CGI and DNA methylation at gene promoters is well understood, the significance of CGIs at 3’ untranslated regions (3’UTRs) remains largely unexplored. In this study, we characterised CGIs located within 3’UTRs and investigated their role in gene regulation. Here, we show that around 3% of CGIs (909 CGIs) are exclusively located at 3’UTR and are not associated with any nearby promoters. Importantly, 3’UTR CGIs are highly conserved, with a conservation score even higher than promoter CGI implying their importance. In contrast to promoter CGIs which are predominantly unmethylated, 3’UTR CGIs are often methylated. Genes with 3’UTR CGIs are associated with several different cancers and cancer-related signalling pathways. Moreover, 3’UTR CGIs are differentially methylated in cancers, with changes in methylation associated with changes in gene expression. Together this data suggests the importance of 3’UTR CGIs and their methylation in gene expression regulation in cancer.

## Background

DNA methylation is a major epigenetic process that regulates gene expression, tissue differentiation, and development^1^. The majority of DNA methylation occurs at CpG dinucleotides, where cytosine is followed by a guanine nucleotide^1^. However, regions with a high density of CpG dinucleotides, and a higher-than-average GC content are called CpG islands (CGI)^2^ and remain protected from methylation. CGIs commonly overlap promoter and enhancer regions, and more than 50% of human genes initiate transcription at regions containing CGIs^3,4^.

Promoter CGIs have been shown to play a significant role in regulating gene expression. It is thought that promoter CGIs are actively protected from methylation to promote open chromatin structure and increase the accessibility of DNA. This facilitates transcription factor and RNA polymerase binding, leading to increased gene expression. Unmethylated promoter CGIs are also associated with the transcription-initiating histone modification H3K4me3^5^. H3K4me3 prevents methylation at promoter CGIs by impairing the binding and activity of *de novo* DNA methyltransferases (DNMTs)^5^. In cancer, DNA methylation has been identified as a key epigenetic mechanism that influences gene expression changes favouring cell proliferation. While cancer cells display a genome-wide loss of CpG methylation, locus-specific hypermethylation of promoter CGIs occurs. The methylation of promoter CGIs results in impaired transcription factor binding and the recruitment of repressive methyl-binding protein, thus leading to stable gene silencing^6^.

Although much of the research has focused on promoter CGIs, a significant number of CGIs are also located across the gene body and intergenic regions^7^. Unlike promoter CGIs, intragenic CGIs often become methylated during development and differentiation^8^. Methylation of intragenic CGIs occurs in a tissue-specific manner, suggesting the process is functionally important for gene regulation^8^. Numerous mechanisms on how intragenic CGIs can influence gene expression have been proposed. Intragenic CGIs have been shown to act as alternative promoters, with methylation of the regions impacting the expression of the host genes, antisense RNAs or microRNAs^7^. In addition, intragenic CGIs have been implicated in pre-mRNA processing mechanisms including alternative polyadenylation^9^ and alternative splicing^10^.

The 3’ untranslated region, 3’UTR, is a untranslated non-coding region of DNA at the 3’ end of the coding sequence of a mRNA^11^. 3’UTRs contains *cis*-regulatory sequences that can be targeted by *trans*-acting factors such as RNA binding proteins and microRNAs^12^. These features of 3’UTRs help regulate gene expression through post-transcriptional mechanisms, including controlling mRNA stability, translation and localisation^11^. Variations in 3’UTR sequence and length can therefore impact gene expression. In the human genome there is a high diversity of 3’UTRs, with over 60% of human genes displaying alternative 3’UTR isoforms^13,14^. This high 3’UTR diversity contributes to the transcriptome landscape, and dysregulation of 3’UTR variants can drive development and progression of diseases such as cancer^15,16^.

A couple of studies which have looked at CGI distribution have identified CGIs within 3’UTRs, although the work was not focused on these sites^17,18^. One study investigating a single 3’UTR CGI found methylation at this site can regulate alternative polyadenylation and cleavage. Here, demethylation of DNA enables CTCF binding to an unmethylated 3’UTR CGI. This allows CTCF to recruit the cohesion complex and form chromatin loops which blocks transcriptional elongation and promotes usage of proximal poly(A) sites^19^. In addition, a study looking at 3’UTR of HAVCR2 gene, found 3’UTR methylation is associated with upregulation genes in several cancer cell lines^20^.

Therefore, it appears DNA methylation may play an important role in regulating 3’UTR diversity, and CGIs located within 3’UTR could be functionally important. In this study, we aimed, for the first time, to characterise CGIs located within 3’UTRs and investigated their role in gene regulation in cancer.

## Materials and Methods

### Identification of 3’UTR CGI

CpG island track^21^ was downloaded from UCSC genome browse (hg38). In this data set CGIs are identified by having a GC content of 50% or greater, a length greater than 200 bp and a ratio greater than 0.6 of observed number of CG dinucleotides to the expected number on the basis of the number of Gs and Cs in the segment. To identify the location of the CGIs in the genome we used the annotatePeak function in the ChIPseeker^22^ package in R. The genomic annotation priority was promoter, 3’UTR, 5’UTR, intron, downstream and intergenic, and the TSS region was set at - 3000 to 3000 kb. The co-ordinates for CGIs which overlapped 3’UTRs and promoter regions were used for downstream analysis. We also generated a list of random 3’UTR coordinates of the same size and distribution that did not overlap a 3’UTR CGI using bedtools^23^ shuffleBed function. To identify genes which have 3’UTR CGIs we used bedtools^23^ intersect function using CGI coordinate file and gencodeV43 annotation file^24^.

### Characterisation of 3’UTR CGI

To assess conservation, we used the 100 vertebrates Basewise Conservation by PhyloP^25^ (phyloP100way) score from UCSC genome browser. To visualise the conservation score at 3’UTR CGIs, promoter CGIs and random 3’UTR coordinates, the phyloP100way score was used to plot heatmaps and metagene plots using deepTools^26^. To identify promoter CGIs which lie within genes with 3’UTR genes, we visualised the intersection between promoter CGIs and 3’UTR CGI gene lists using the UpSet^27^ function of the Bioconductor package ComplexHeatmap^28^ in R. The CpG island track from UCSC genome browser provides information on CGI length, percentage of CGI containing CpG and GC, and observed:expected CpG. The data was plotted using Prism. To identify the nearest TSS to 3’UTR CGI, annotatePeaks.pl function in HOMER^29^ was used. The Distance to TSS Nearest PromoterID was plotted. Processed ChIP-seq data was accessed from the ENCODE database (RNAPII HEPG2 [GEO:GSM935603], RNAPII MCF7 [GEO:GSM1006865], RNAPII HELA [GEO:GSM822273], H3K4me3 HEPG2 [GEO:GSE96248], H3K4me3 MCF7 [GEO:GSM945269], H3K4me3 HELA [GEO:GSE96127], H3K36me3 HEPG2 [GEO:GSM733685], H3K36me3 MCF7 [GEO:GSE174945], H3K36me3 HELA [GEO:GSM945230]). Metagene plots using ChIP data were generated using deepTools^26^.

### Characterisation of 3’UTR CGI genes

Tau Index V8^30^ was used to assess tissue specificity of genes with 3’UTR CGIs compared with a list of housekeeping genes. Gene ontology was performed using the Bioconductor package clusterProfiler^31^ in R. Pathway analysis as performed using g:Profiler^32^.

### Analysis of expression and methylation data from TCGA

Breast Invasive Carcinoma (BRCA) and hepatocellular carcinoma (LIHC) expression, 450K methylation array data and clinical data were downloaded from The Cancer Genome Atlas (TCGA)^33^ using Bioconductor package TCGABiolinks^34^ in R. Differential gene expression analysis was performed by DESeq2^35^ in R. Differential analysis of CpG probes was performed using the Bioconductor package Limma^36^ in R. For each CpG probe, a linear model was fit to the intensity values across all samples using lmFit function, and the model statistics and p-values of differential methylation were computed using the eBayes function. The top 10 differentially methylated probes in 3’UTR CGIs were plotted using the plotCpg function. The Bioconductor package DMRcate^37^ was used to identify and analyse DMRs. DMRcate was used at a bandwith of 1000 nucleotides (lambda = 1000) with the scaling factor bandit set at 2 (C = 2) which are the recommended parameters for array data. DMR regions were plotted using the Bioconductor package GViz^38^ in R. CpG Methylation by Methyl 450K Bead Arrays from ENCODE/HAIB for cell lines HMEC, MCF7, Hepatocytes and Hepg2 were visualised using UCSC genome browser.

### Statistical analysis

Statistical analysis was performed using GraphPad Prism or as a part of the computational tools used. To measure statistical significance, Brown-Forsythe and Welch ANOVA tests were used (Fig 1E, Fig 2A), or paired T-test (Fig 4Eiii) were used to calculate P-value. P-values are provided in the figure legends or indicated on the figures. P-values of <0.05 were considered significant.

**Figure 1:**
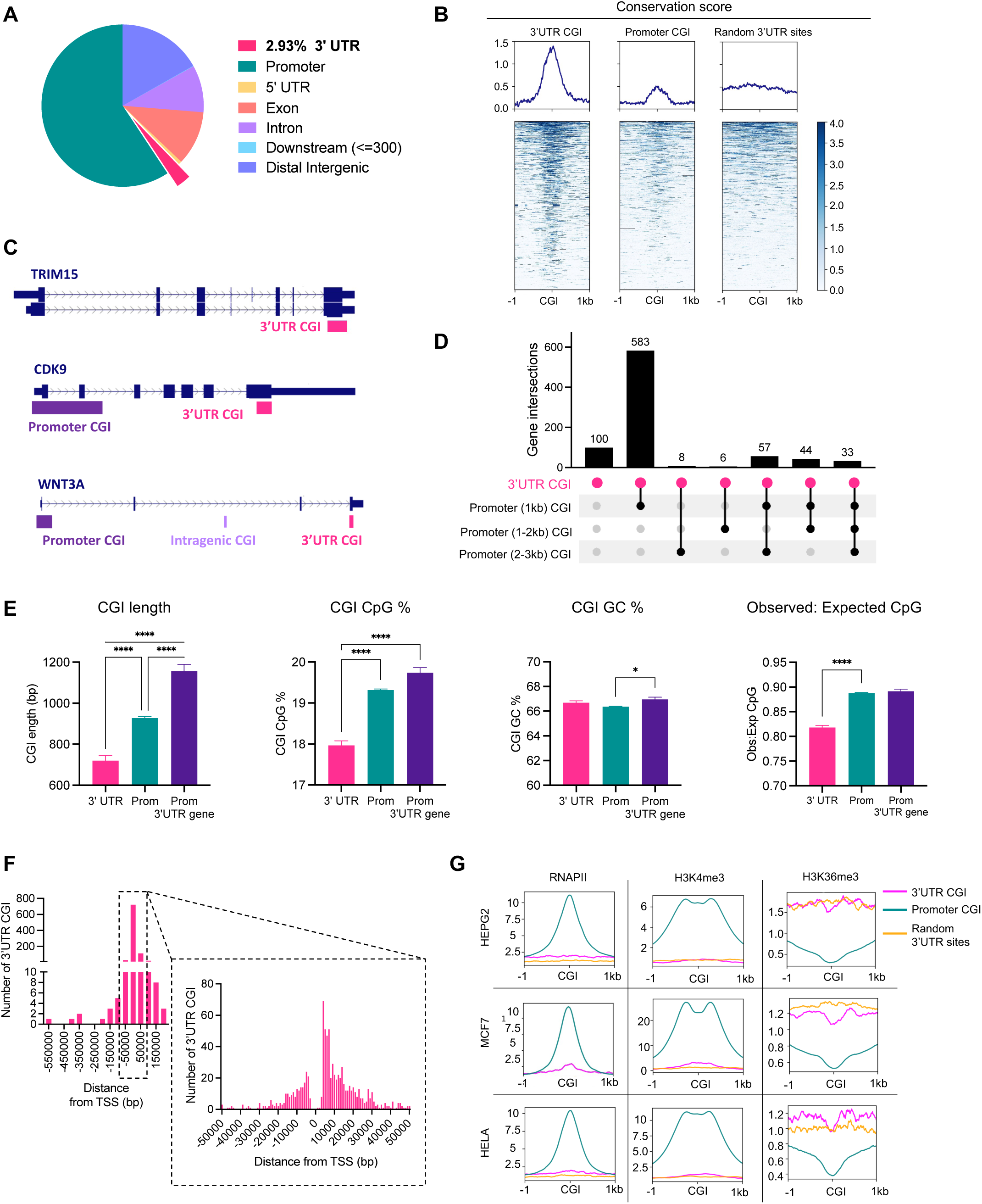
Characterization of 3’UTR CpG islands. A) A pie chart showing the genome distribution of CpG islands (CGIs). B) Metagene plot and heatmaps showing the conservation (PhyloP100way) score of 3’UTR CGIs, promoter CGIs (subset of equal number to 3’TR CGIs) and random 3’UTR sites. C) Example of genes containing a 3’UTR CGI, with presence of promoter CGI and intragenic CGI highlighted. D) UpSet diagram showing the number of genes with 3’UTR CGIs which contain promoter CGIs. E) Bar plots showing CGI length, percentage of CpG withing the CGI, percentage of GC within the CGI and the observed to expected CpG ratio within the CGI. Values represent the mean ± standard deviation. ∗p < 0.05; ∗∗∗∗p < 0.0001 (Brown-Forsythe and Welch ANOVA test). F) A bar plot showing the distance of 3’UTR CGIs from known transcriptional start site (TSS), with a zoomed in bar plot showing number of 3’UTR CGIs located -50000 to 50000 bp from the nearest TSS. G) Metagene plots showing the enrichment of RNA polymerase II (RNAPII), H3K4me3, and H3K36me3 in 3’UTR CGI, promoter CGI and random 3’UTR regions in HEPG2, MCF7 and HELA cell lines.

**Figure 2:**
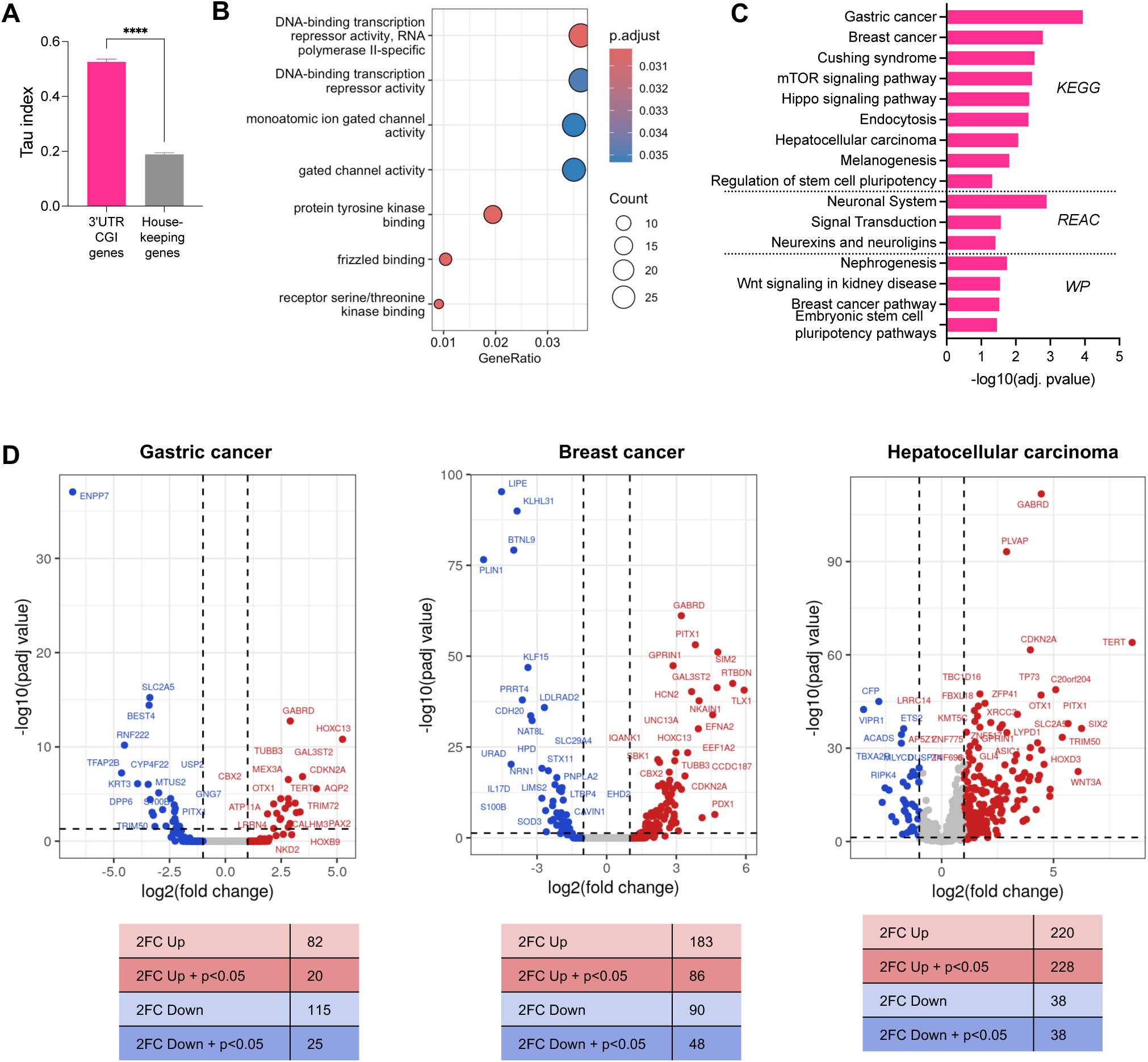
Characterization of genes with 3’UTR CpG islands. A) A bar plots showing the tau index of genes with 3’UTR CGI and list of housekeeping genes. Values represent the mean ± standard deviation. ∗p < 0.05 (Brown-Forsythe and Welch ANOVA test). B) A dot plot of gene ontology enrichment terms for genes with 3’UTR CGIs. The diameter indicates the number of genes overlapping the gene ontology term and the colour indicates the enrichment P-value. C) A bar plot showing enriched pathways in the KEGG, Reactome and WP database for genes with 3’UTR CGIs. D) Volcano plot showing differential expression of 3’UTR CGI genes in gastric cancer, breast cancer, and hepatocellular carcinoma tumour tissue compared to normal tissue. Data from TCGA used. Red dots indicate genes with a 2-fold change upregulation, and blue dots indicated genes with a 2-fold change downregulation.

## Results

### CGIs in 3’UTR regions are more conserved than promoter CGIs

We first sought to characterise CGI in 3’UTR regions and examine them in comparison to promoter CGIs. We identified 909 CGIs (∼3% of all CGIs) that exclusively reside in 3’UTR regions with no promoter region in the vicinity (**Fig 1A**). To assess functional importance of 3’UTR CGIs, we calculated the conservation score at these CGIs. Conservation of genomic element indicate their functional importance, as these regions are under positive selection during evolution. Therefore, if 3’ UTR CGIs are functionally important, they should display conservation score at least comparable to promoter CGIs which are well-known to play important role in gene expression^39^. Surprisingly, our analyses shows that 3’UTR CGIs are not only highly conserved but display a conservation score even higher than promoter CGI (**Fig 1B**). Previous studies have shown highly conserved regulatory elements involved in transcriptional termination and post-transcriptional regulation are present in 3’ UTRs^40^. Therefore, the high conservation score of 3’UTR CGIs might reflect these regulatory elements present in 3’UTR rather than CGIs themselves. To verify this, we compared the conservation of 3’ UTR CGIs to conservation of a control set consisting of 3’UTR regions of similar size. A comparison between 3’UTR CGI and control set shows that CGI at 3’UTR are much more conserved than non-CGI 3’ UTR regions (**Fig 1B**). The high conservation at 3’UTR CGI suggests possible function significance for these CGIs.

Most genes contain CGIs within their promoter regions. We wanted to assess if genes with 3’UTR CGIs also contain promoter CGIs. Broadly, 3’UTR CGI genes can be divided into three categories - with 3’UTR CGIs alone, with promoter and 3’UTR CGIs, or contain promoter and 3’UTR CGIs as well as other intragenic CGIs (**Fig 1C**). The UpSet plot of 3’UTR CGI containing genes show that only a small number of genes (100 genes) have CGI only at the 3’ end. A large majority of genes (731 genes) also contain promoter CGIs (**Fig 1D**). Moreover, majority of these promoter CGIs are within 1kb of the transcriptional start site.

The features of CGIs can influence their activity^41^. Therefore, we compared the composition of 3’UTR CGIs with composition of promoter CGIs (**Fig 1E**). The comparison showed that 3’UTR CGIs are significantly shorter than promoter CGIs, with a lower amount of CpGs. This indicates that 3’ UTR CGIs are lesser in strength as compared to promoter CGIs. Most genes with 3’UTR CGIs also contain promoter CGIs. It is possible that this subset of genes contain promoter CGIs which have distinct characteristics, such as being longer in length, which could influence the expression profile of these genes^41^. Therefore, we wanted to see if these promoter CGIs at genes also associated with CGIs at 3’UTR are different from other promoter CGIs where the gene is not associated with 3’UTR CGI. We found that 3’UTR-associated promoter CGIs are longer in length to other promoter CGIs, although the level of CpG is comparable. A study has shown that longer promoter CGIs contain more RNA polymerase II (Polr2a) binding sites and transcriptional start sites^41^. This suggests that genes with CGIs at both promoter and 3’UTRs might have different transcriptional status compared to genes with CGIs only atpromoters.

CGIs are commonly found near transcriptional start sites (TSS). One initial explanation for the presence of 3’UTR CGIs is that they act as TSSs and initiate gene expression of a near-by or antisense transcript. We assessed the distance between 3’UTR CGIs and the nearest TSS (**Fig 1F**). All 3’UTR CGIs are located at least 3000bp away from an annotated TSS, with the majority of TSS located at least 5000bp away from a 3’UTR CGI, suggesting 3’UTR CGIs do not act to promote transcription initiation. To further explore the potential role of 3’UTR CGIs as sites of transcription, we analysed RNA polymerase II (RNAPII) binding and enrichment of the transcription-initiating histone modification H3K4me3 in several cell lines (**Fig 1G**). In 3’UTR CGIs, there is little evidence of RNAPII or a H3K4me3, while promoter CGIs are enriched for both. We also looked at binding of the histone H3K36me3 which is associated with transcription elongation and is correlated with DNMT3B-mediated methylation^42^. We see low levels of H3K36me3 binding in promoter CGIs, which correlates with these sites being predominately sites of transcription initiation rather than transcription elongation. At 3’UTR CGI sites, H3K36me3 shows higher enrichment than at the promoter CGIs but it is comparable to a control set of non-CGI 3’UTR regions. Together, the data shows 3’UTR CGIs are distinct to promoter CGIs, with no evidence of 3’UTR CGIs acting as alternative TSS.

### Genes with CpG islands in 3’UTRs are associated with cancer and cancer-related signalling

CGIs are often associated with housekeeping genes that are constitutively expressed in majority of cells in tissue-non-specific manner. Given that 3’UTR CGIs are highly conserved and mostly associated with a promoter CGIs, it is possible they act as an extra layer of regulation for housekeeping genes. To verify this, we measured the tissue specificity of genes with 3’UTR CGIs and found 3’UTR CGI genes have a significantly higher tissue specificity score than housekeeping genes (**Fig 2A**). This suggests genes with 3’UTR CGI may have tissue-specific roles.

To further investigate the role of 3’UTR CGIs, we checked if genes with 3’UTR CGIs are involved in specific functional pathways by carrying out gene ontology analyses. Enriched molecular functions in 3’UTR CGI genes included transcription regulation and transcription factor binding (**Fig 2B**) indicating tissue-specific expression of these genes. Interestingly, pathway analysis showed terms related to cancer are enriched in genes with 3’UTR CGIs (**Fig 2C**). Terms included gastric, breast and hepatocellular carcinoma as well as the mTOR and Hippo signalling which are key pathways dysregulated in cancer^43,44^. Enrichment is also seen in processes which have been shown to be disrupted in tumour initiation and progression, including endocytosis^45^, melanogenesis^46^, and neuronal signalling^47^.

We also assessed if the expression levels of genes with 3’UTR CGIs change in cancer. We looked at gastric, breast and hepatocellular carcinoma as they had been highlighted in pathway analysis. Using gene expression data from patient samples (available via TCGA) we found 3’UTR CGI genes are differentially expressed in cancer, with the expression profiles varying upon cancer types (**Fig 2D**). The work further suggests a role for 3’UTR CGIs in regulating genes involved in cancer.

### 3’UTR CpG islands are differentially methylated in cancer

Cancer cells are characterized by genome-wide hypomethylation and hypermethylation of CGIs at promoters^6^. Given that 3’ UTR CGI genes are differentially expressed in cancer and associated with key cancer pathways, we hypothesised that like promoter CGIs, 3’ UTR CGIs are differentially methylated in cancer. For this analysis we focused on breast cancer (BRCA) and hepatocellular carcinoma (LIHC) where methylation data from matched normal tissue was available to assess cancer-related differential methylation.

Using methylation array data, we assessed the methylation value of individual CpG probes which represent a single-CpG-site (**Fig 3A**). In both normal and tumour samples, 3’UTR CGIs are highly methylated, with a methylation score closer to 1 (**Fig 3B**). In comparison, promoter CGIs remain unmethylated in both conditions. There was no clear difference in overall methylation of CpGs between normal and tumour samples for both 3’UTR and promoter CGIs perhaps because the changes in methylation occur at subset of CpG sites depending on cancer type. To check if this is the case, we identified methylation data from BRCA and LIHC. It is clear from these analyses, that in both cancers’ methylation changes occur at 3’UTR CGIs (**Fig 3C, D**). However, overall methylation pattern is cancer specific. In BRCA samples, 3’UTR CGIs are more likely to become hypermethylated, whereas in LIHC, both hyper-and hypo-methylation of 3’UTR CGIs can be seen. To further investigate the impact of 3’UTR CGI methylation may have in cancer, we identified the top 10 significantly differentially methylated CpG sites in 3’UTR CGI regions and associated genes (**Fig 3E**). Interestingly, many of the genes identified with having differentially methylated 3’UTR CGIs had previously been associated with BRCA, LIHC or other cancers (**Suppl. Table 1**), further suggesting a functional role for 3’UTR CGIs in cancer.

**Figure 3:**
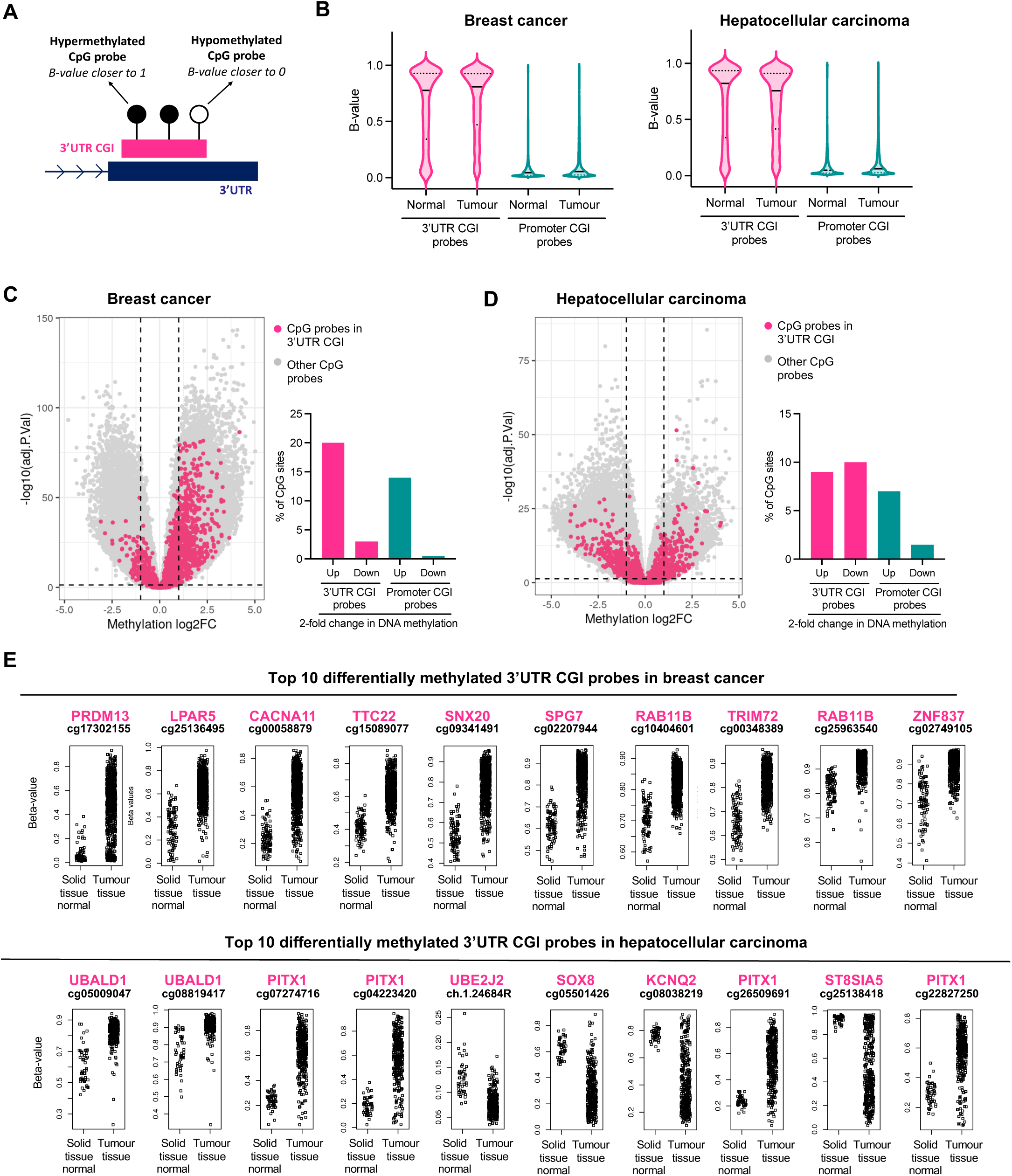
Differential methylation of 3’UTR CpG islands in cancer. A) A schematic explaining CpG probes assessed in methylation array data. B) Volcano plots showing methylation B-values for CpG probes in 3’UTR CGIs and promoter CGIs in normal and tumour samples in breast cancer and hepatocellular carcinoma. A volcano plot showing differential methylation of CpG probes in C) breast cancer and D) hepatocellular carcinoma tumour tissue compared to normal tissue. Pink dots indicate CpG probes in 3’UTR CGI. A bar plot showing the percentage of CpG probes which are methylated and unmethylated in breast cancer and hepatocellular carcinoma tumour samples. E) Scatterplot dot plots showing the methylation beta values of the top 10 differentially methylated CpG probes in 3’UTR CGIs in breast cancer and hepatocellular carcinoma.

We next focused on differentially methylated regions (DMRs) which are genomic regions consisting of several differentially methylated CpG (**Fig 4A**). Compared to individual differentially methylated CpG sites, DMRs are biologically more relevant and likely to be associated with transcriptional changes^48^. In both BRCA and LIHC, DMRs were found to be located within 3’UTR CGIs (**Fig 4B**). Genes with 3’UTR CGI DMRs had previously been associated with BRCA, LIHC or other cancer (**Suppl. Table 2**). In BRCA, the majority of 3’UTR CGIs were hypermethylated in tumour tissue. In LIHC, 3’UTR CGIs were both hyper-and hyo-methylated in tumour tissue. As changes in promoter methylation can influence gene expression profiles, we assessed gene expression changes in genes which contained a 3’UTR CGI DMR, but did not have differential methylation in their promoter CGIs (**Fig 4C**). This way we could separate the effect of promoter methylation from methylation at 3’ UTR. Here, we saw a trend for genes with a hypermethylated DMR within their 3’UTR CGI to have increased expression in BRCA tumour tissue, while genes with hypomethylated DMR in the 3’UTR CGI are more likely to have decreased expression in tumour tissue (**Fig 4D**). In LIHC, a similar trend in genes with hypermethylated 3’UTR CG DMRs having increased gene expression in LIHC tumour tissue was seen. However, hypomethylated 3’UTR CGI DMRs did not show any correlation with expression (**Fig 4D**).

**Figure 4:**
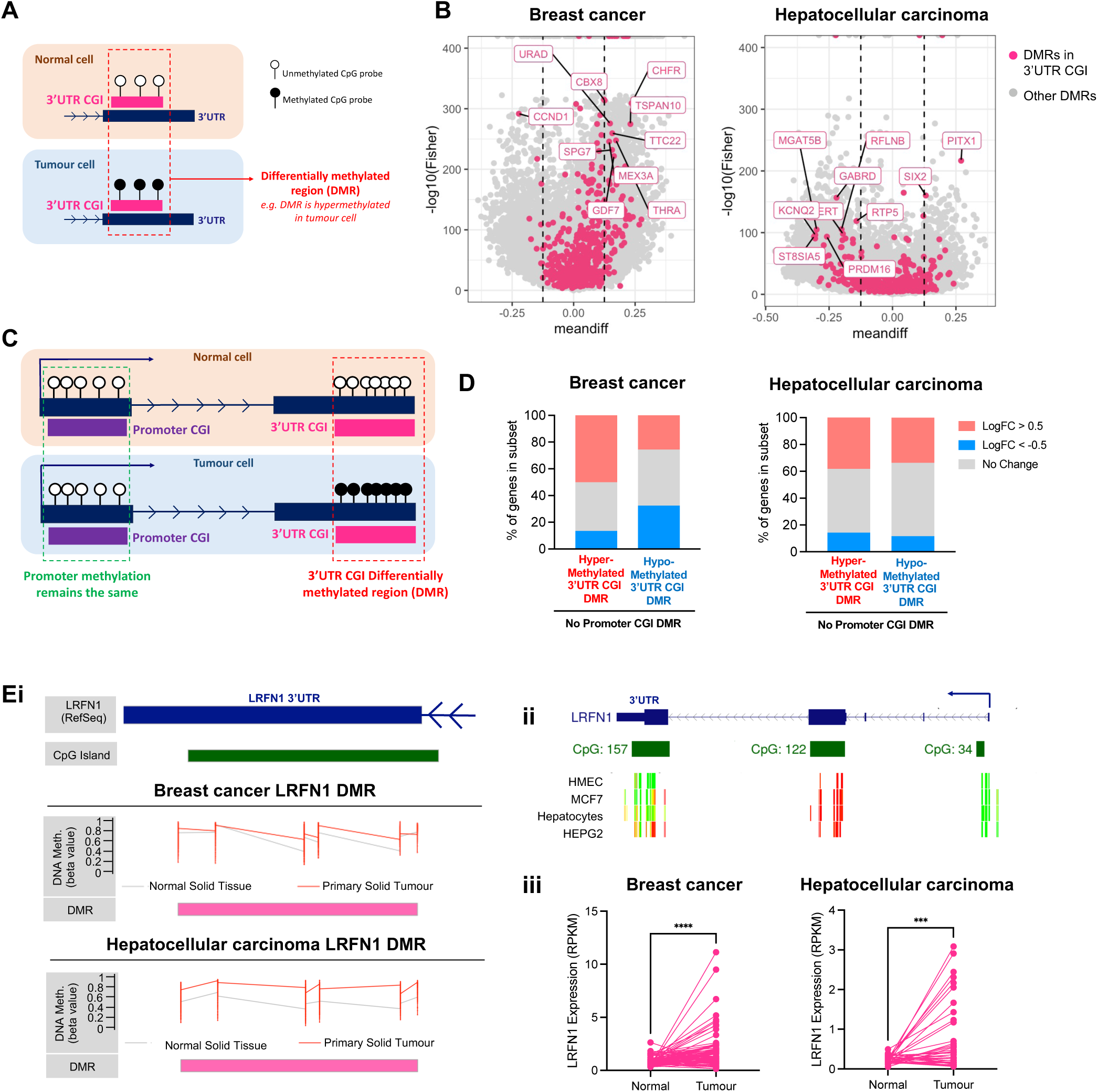
Identification of differentially methylated regions in 3’UTR CpG islands in cancer. A) A schematic explaining differentially methylated regions (DMR). B) Volcano plot showing DMRs (differentially methylated in tumour compared to cancer) in breast cancer and hepatocellular carcinoma. Top 10 DMRS are highlighted (based on meandiff > 0.125 and ranked based on P-value). Pink dots indicate 3’UTR CGI DMRs. C) A schematic showing an example of a gene with a DMR in the 3’UTR CGI but no DMR in the promoter CGI region. D) Bar plots showing the percentage of genes with a 3’UTR DMR but no promoter CGI DMR which are upregulated (red, LogFC >0.5) and downregulated (blue, LogFC < -0.5) in breast cancer and hepatocellular carcinoma. Ei) Figure showing methylated DMR in the gene LRFN1 in breast cancer and hepatocellular carcinoma dataset. Eii) UCSC track diagram showing methylation regions across LRFN in HMEC (primary mammary cells), MCF-7 (breast cancer cells), hepatocytes (control hepatocyte cells) and HEPG2 (hepatocellular carcinoma cells). Red lines indicated methylated regions; green lines indicate unmethylated regions. Eii) Scatter plot showing expression of LRFN1 in paired patient data from breast cancer (n=77) and hepatocellular carcinoma dataset (n=40). ∗∗∗∗p < 0.0001 (paired T-test).

A good example of link between hypermethylation of CGIs with expression is LRFN1 gene. This gene shows hypermethylated DMR in the 3’UTR CGI region but no change in methylation in promoter CGIs (**Fig Ei**). Hypermethylation is also seen in LRFN1 3’UTR in MCF7 cells (breast cancer cell) and HEPG2 (hepatocellular carcinoma cell line) compared to control cell lines (**Fig Eii**). We assessed expression of LRFN1 in paired patient data, which further confirmed LRFN1 increases in expression in BRCA and LIHC compared to patient-matched control tissue (**Fig Eiii**).

## Discussion

The majority of research investigating CpG islands have focused on promoter regions, with ∼70% of gene promoters identified to be within CGIs^1^. While a couple of studies investigating CGI genomic distribution have identified CGIs within 3’UTRs, however, these studies have not explored these sites in a high degree of detail^17,18^. Understanding how site-specific DNA methylation contributes to gene expression regulation remains an important research question. With 3’UTRs being sites of post-transcriptional gene regulation, methylation changes occurring at 3’UTR CGIs can potentially influence gene expression and offer additional mechanism of regulation to explain gene expression changes that cannot be explained merely through promoter methylation.

In this study we began by identifying and characterising 3’UTR CGIs. Even though only ∼3% CGIs (**Fig 1A**) exclusively reside within 3’UTR regions, it represents a significant number of genes (831) large proportion of which are identified to be tissue-specific genes involved in cancer pathways (**Fig 2**). The importance of 3’ UTR CGIs is also reflected in our observation that 3’UTR CGIs are much highly conserved compared to promoter CGIs as well as other 3’UTR regions. High conservation of 3’UTR CGIs has not been reported before but reflects that 3’UTR CGIs are of equal, if not more, importance as compared to promoter CGIs.

We further aimed to elucidate the role of 3’UTR CGIs. Previous reports indicate that RNA polymerase has affinity for CGIs^4^. In addition, presence of antisense transcripts is a common phenomenon in the human genome, and intragenic CGIs have previously been shown to act as alternative TSS^7^. Together this made us speculate that 3’ UTR CGIs might be sites of transcription. However, our data suggests that 3’UTR CGIs were neither located near annotated TSS nor there was any evidence of RNAPII and H3K4me3 enrichment (**Fig 1G**).

Aberrant methylation of promoter CGIs is a key contributor to cancer initiation and progression^49^. Therefore, it is likely that 3’ UTR CGIs are also subjected to aberrant methylation during cancer. An examination of methylation status of 3’UTR CGIs indicates that these CGIs have higher level of methylation as compared to promoter CGIs which are protected from methylation (**Fig 3B**). It is often hypothesised that human genome is relatively devoid of CpG dinucleotides because methylated cytosines are prone to mutagenesis leading to their dilution over time^50^. However, CGIs are protected from methylation, leading to their conservation and high CpG density. On this background, our observation that 3’UTR CGIs are more likely to be methylated is significant. It might explain their smaller size and lower density. However, the observation that they are highly conserved points not only to their significance but also raises interesting questions regarding the mechanisms behind their preservation and the impact of their demethylation on gene regulation.

3’UTR CGIs were differentially methylated in the two cancers that we studied (**Fig 3**, **4**). Interestingly, genes with differentially methylated CpG sites or DMRs within 3’UTR CGIs had previously been associated with cancer, further supporting the concept that 3’UTR CGIs play a role in gene regulation in cancer. However, overall methylation pattern of 3’ UTR CGIs was cancer specific, highlighting different mechanisms of oncogenesis. Previous pan-cancer studies have made similar observations, showing distinct DNA methylation patterns in different cancers^51–53^. Similar to our data, breast cancer has been shown to contain a high proportion of hypermethylation, while hepatocellular carcinoma has more evidence of differential hypomethylation^51,53^. Therefore, it is unsurprising that we see differential patterns of 3’UTR CGIs in breast cancer and hepatocellular carcinoma.

As DMRs are biologically more relevant, we further looked at genes with 3’UTR CGI DMRs. Particularly, we focused on genes which did not contain a DMR in a promoter CGI as changes as promoter methylation is known to influence gene expression. Here, we found a subset of genes that displayed a methylated 3’UTR CGI showed increased gene expression (**Fig 4D**). In contrast genes with a hypomethylated DMR in the 3’UTR CGI displayed decreased expression in tumour tissue, especially evident in breast cancer data. The correlation between increased methylation at the 3’UTR CGI with increased transcription can be a consequence rather than the cause^54^. However, a closer look at selected genes, e.g. LRFN1 we see gene body methylation in both control and tumour samples. The 3’UTR CGI regions is the only region differentially methylated in cancer (**Fig 4E**). Moreover, our observation that 3’UTR CGIs are found in tissue-specific genes rather than ubiquitously expressed housekeeping genes indicates that 3’UTR CGI methylation might provide additional layer of specificity to gene expression. Previously published example of immune checkpoint gene HAVCR2 which displays increased 3’UTR DNA methylation and increased expression upon T cell activation further supports our observation that 3’UTR CGI hypermethylation can lead to increased gene expression.^20^.

However, the question remains how might 3’UTR CGIs and their methylation influence gene expression. It is possible CGIs might influence 3’ UTR function. For example, methylation of 3’ UTR CGIs can regulate polyadenylation signals (poly-A tail). Most genes contain multiple methylation signal. Usage of these signals impacts 3’UTR length and therefore post-transcriptional regulation including mRNA stability, translation and localisation^11^. A previous study has suggested that 3’UTR methylation can promot the use of proximal polyA sites through CTCF-mediated chromatin looping^19^. 3’UTR methylation could also influence 3’UTR length through regulation of RNA Pol II speed. Altered rates of RNA Pol II can impact splicing, polyadenylation and the production of alternative splice variants^10^. Intragenic methylation has been shown to lead to the slowing of RNA Pol II, which in this example allowed for weak splice sites to be recognised^10^. It is possible that a similar mechanism may occur at 3’UTR CGI sites, with methylation of 3’UTR CGIs resulting in slowing of RNA Pol II and therefore influencing choice of poly(A) sites. This would result in shorter 3’UTR associated increased stability of mRNAs. Alternatively, 3’UTR functions are mediated by the binding of proteins to the 3’UTR. Many transcription factor bindings sites have been found to be GC rich, with CGIs acting to enhance protein binding^4^. Furthermore, CGI methylation can block the binding of transcription factors in these regions. Therefore, it is possible that 3’UTR CGIs act as binding sites for proteins involved in splicing and stability, with differential methylation of 3’UTR CGIs resulting in changes to protein binding and thus gene expression.

In summary, this study characterises 3’UTR CGIs for the first time and identifies them as conserved features of tissue-specific genes. The work indicates that methylation of these 3’UTR CGIs can regulate expression of genes in cancer pathway. The work highlights the importance of of DNA methylation in sites other than promoter can be also highly important in gene regulation.

---

**Suppl. table 1:**
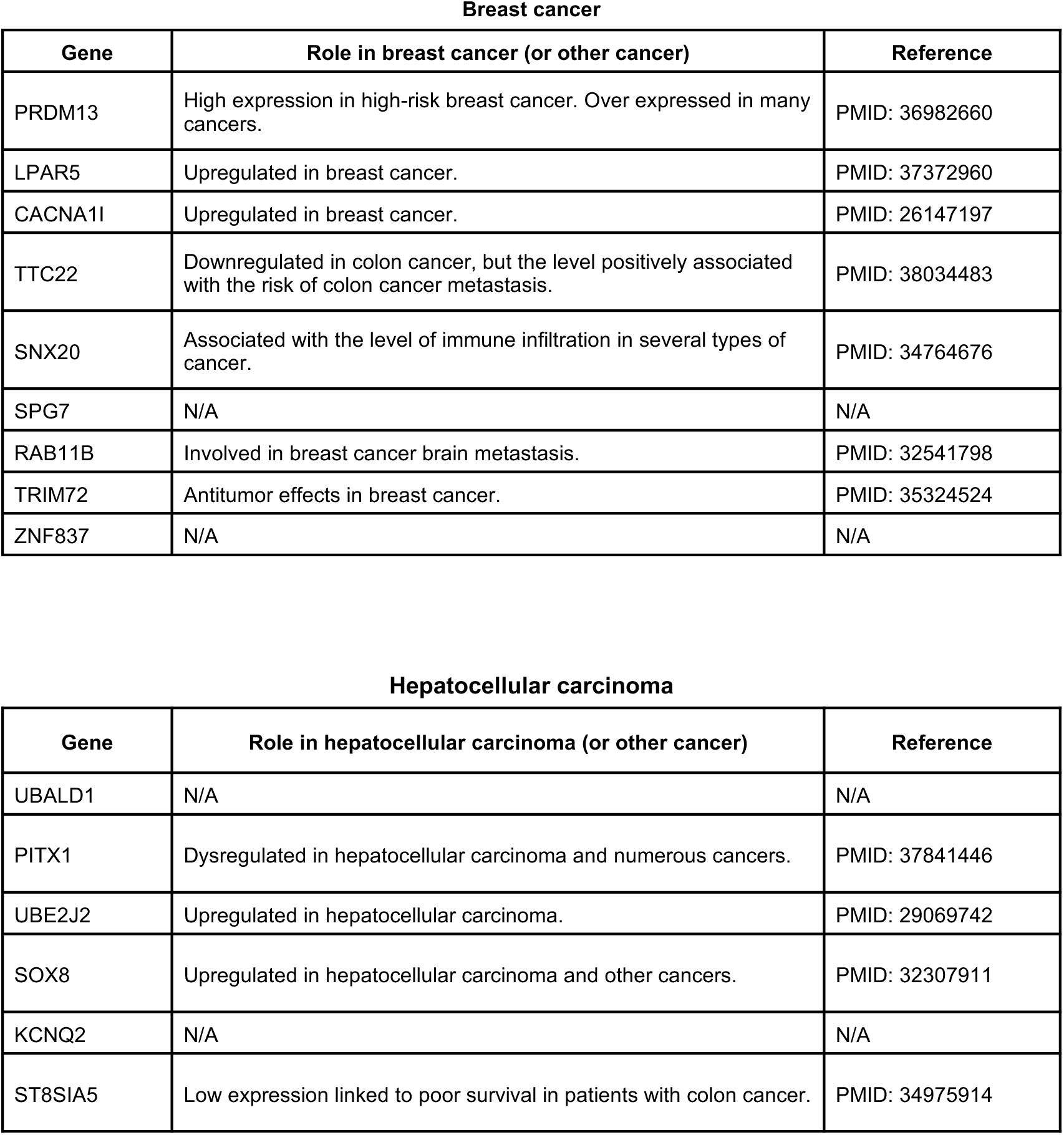
Role of genes associated with top 10 differentially methylated 3’UTR CGI probes in BRCA and LIHC. Genes which contain top 10 differentially methylated 3’UTR CGI probe in either breast cancer or hepatocellular carcinoma, with brief description of known role in cancer.

Breast cancer
| Gene | Role in breast cancer (or other cancer) | Reference |
| --- | --- | --- |
| PRDM13 | High expression in high-risk breast cancer. Over expressed in many cancers. | PMID: 36982660 |
| LPAR5 | Upregulated in breast cancer. | PMID: 37372960 |
| CACNA1I | Upregulated in breast cancer. | PMID: 26147197 |
| TTC22 | Downregulated in colon cancer, but the level positively associated with the risk of colon cancer metastasis. | PMID: 38034483 |
| SNX20 | Associated with the level of immune infiltration in several types of cancer. | PMID: 34764676 |
| SPG7 | N/A | N/A |
| RAB11B | Involved in breast cancer brain metastasis. | PMID: 32541798 |
| TRIM72 | Antitumor effects in breast cancer. | PMID: 35324524 |
| ZNF837 | N/A | N/A |

**Suppl. table 1: Role of genes associated with top 10 differentially methylated 3'UTR CGI probes in BRCA and LIHC.**
| Gene | Role in hepatocellular carcinoma (or other cancer) | Reference |
| --- | --- | --- |
| UBALD1 | N/A | N/A |
| PITX1 | Dysregulated in hepatocellular carcinoma and numerous cancers. | PMID: 37841446 |
| UBE2J2 | Upregulated in hepatocellular carcinoma. | PMID: 29069742 |
| SOX8 | Upregulated in hepatocellular carcinoma and other cancers. | PMID: 32307911 |
| KCNQ2 | N/A | N/A |
| ST8SIA5 | Low expression linked to poor survival in patients with colon cancer. | PMID: 34975914 |

**Suppl. table 2:**
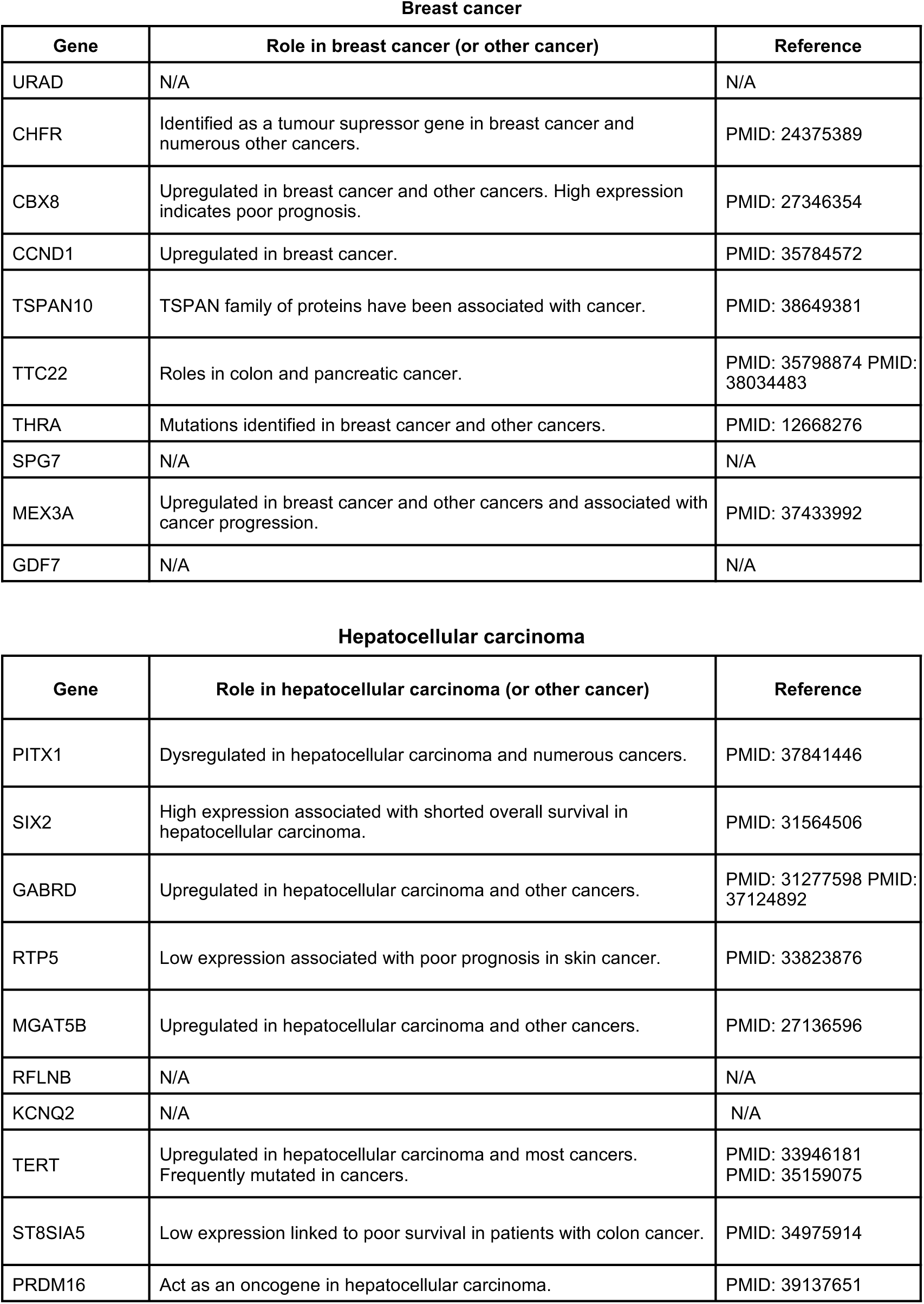
Role of genes associated with top 10 3’UTR CGI differentially methylated regions in BRCA and LIHC. Genes which contain top 10 3’UTR CGI differentially methylated regions (DMRs) in either breast cancer or hepatocellular carcinoma, with brief description of known role in cancer.

Breast cancer
| Gene | Role in breast cancer (or other cancer) | Reference |
| --- | --- | --- |
| URAD | N/A | N/A |
| CHFR | Identified as a tumour suppressor gene in breast cancer and numerous other cancers. | PMID: 24375389 |
| CBX8 | Upregulated in breast cancer and other cancers. High expression indicates poor prognosis. | PMID: 27346354 |
| CCND1 | Upregulated in breast cancer. | PMID: 35784572 |
| TSPAN10 | TSPAN family of proteins have been associated with cancer. | PMID: 38649381 |
| TTC22 | Roles in colon and pancreatic cancer. | PMID: 35798874 PMID: 38034483 |
| THRA | Mutations identified in breast cancer and other cancers. | PMID: 12668276 |
| SPG7 | N/A | N/A |
| MEX3A | Upregulated in breast cancer and other cancers and associated with cancer progression. | PMID: 37433992 |
| GDF7 | N/A | N/A |

**Suppl. table 2: Role of genes associated with top 10 3'UTR CGI differentially methylated regions in BRCA and LIHC.**
| Gene | Role in hepatocellular carcinoma (or other cancer) | Reference |
| --- | --- | --- |
| PITX1 | Dysregulated in hepatocellular carcinoma and numerous cancers. | PMID: 37841446 |
| SIX2 | High expression associated with shortened overall survival in hepatocellular carcinoma. | PMID: 31564506 |
| GABRD | Upregulated in hepatocellular carcinoma and other cancers. | PMID: 31277598 PMID: 37124892 |
| RTP5 | Low expression associated with poor prognosis in skin cancer. | PMID: 33823876 |
| MGAT5B | Upregulated in hepatocellular carcinoma and other cancers. | PMID: 27136596 |
| RFLNB | N/A | N/A |
| KCNQ2 | N/A | N/A |
| TERT | Upregulated in hepatocellular carcinoma and most cancers. Frequently mutated in cancers. | PMID: 33946181 PMID: 35159075 |
| ST8SIA5 | Low expression linked to poor survival in patients with colon cancer. | PMID: 34975914 |
| PRDM16 | Act as an oncogene in hepatocellular carcinoma. | PMID: 39137651 |

